## Supplementary Figures for "A Unified Pipeline for FISH Spatial Transcriptomics"

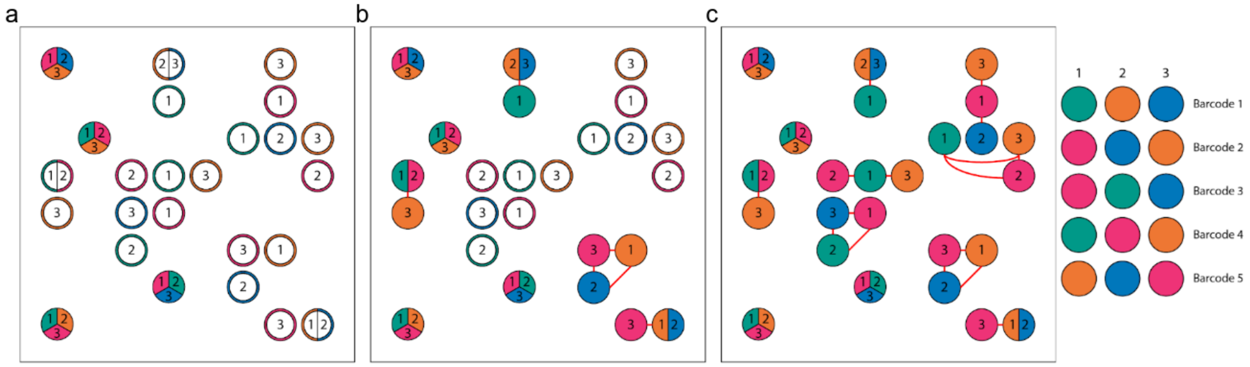


**Figure S1**: **Schematic representation of the results of different seqFISH decoding methods**

Each circle represents a spot in a theoretical seqFISH experiment with the number denoting the imaging round and the color corresponding to the imaging channel. Split spots show spatially co-occurring spots, filled in spots are part of a decodable barcode for the decoding method shown in that panel and red lines connect spots of the same decodable barcode that do not spatially co-occur. The codebook is shown on the far right. Decoding methods shown are **a)** starfish ExactMatch decoder, **b)** starfish NearestNeighbor decoder, and **c)** the custom CheckAll decoder.

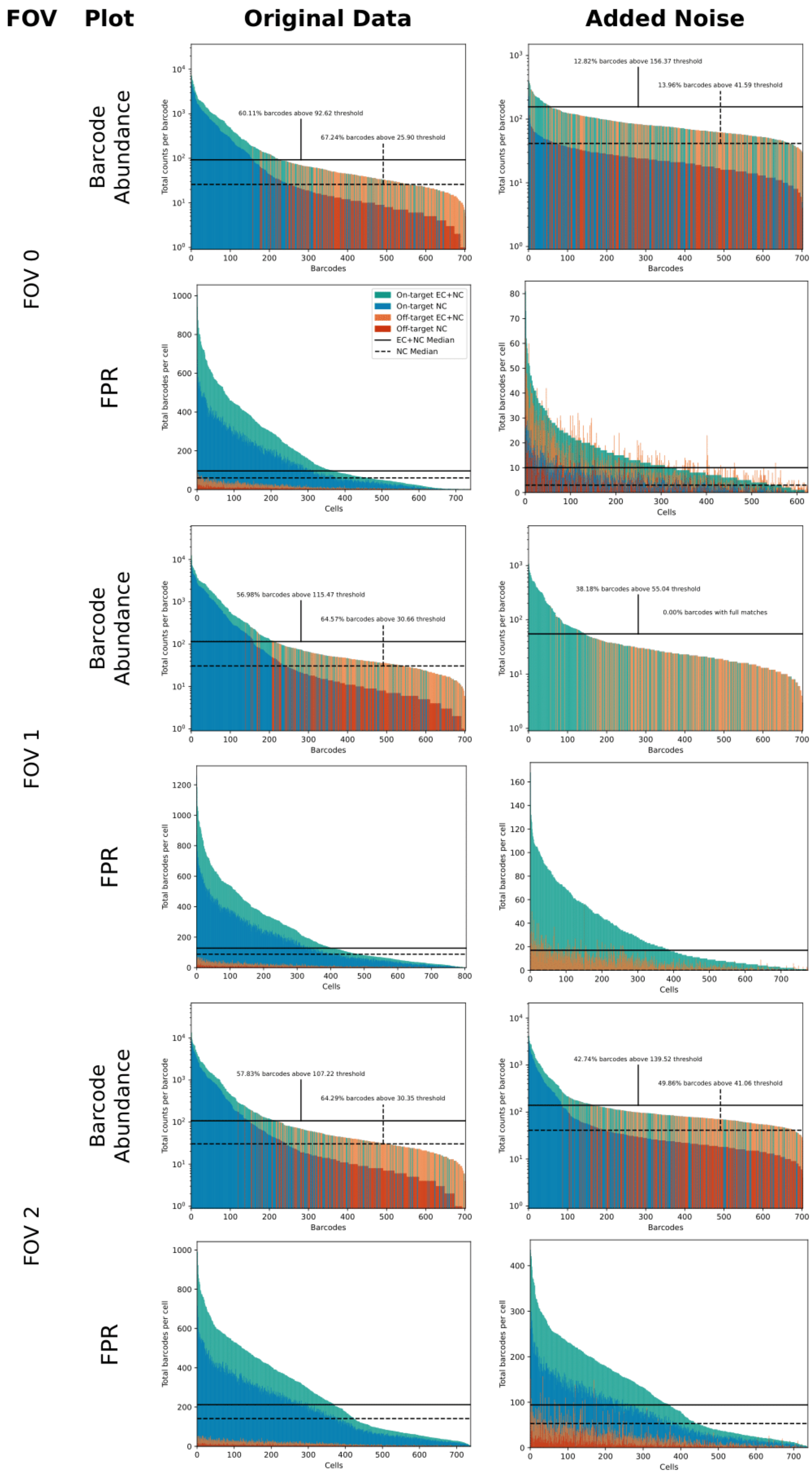
FOV
Plot
Original Data
Added Noise
FOV 0
FOV 1
FOV 2
Barcode Abundance
FPR
60.11% barcodes above 92.62 threshold
67.24% barcodes above 25.90 threshold
12.82% barcodes above 156.37 threshold
13.96% barcodes above 41.59 threshold
56.98% barcodes above 115.47 threshold
64.57% barcodes above 30.66 threshold
38.18% barcodes above 55.04 threshold
0.00% barcodes with full matches
57.83% barcodes above 107.22 threshold
64.29% barcodes above 30.35 threshold
42.74% barcodes above 139.52 threshold
49.86% barcodes above 41.06 threshold
Total counts per barcode
Barcodes
Total barcodes per cell
Cells
On-target EC+NC
On-target NC
Off-target EC+NC
Off-target NC
EC+NC Median
NC Median

**Figure S2**: **Comparison of selected QC metrics on data before and after simulated noise.** Original data taken directly from pipeline output of seqFISH data, “Added Noise” results were obtained by adding noise to the seqFISH images (**Methods**) and then running the same pipeline. EC = error-corrected, NC = non-corrected.

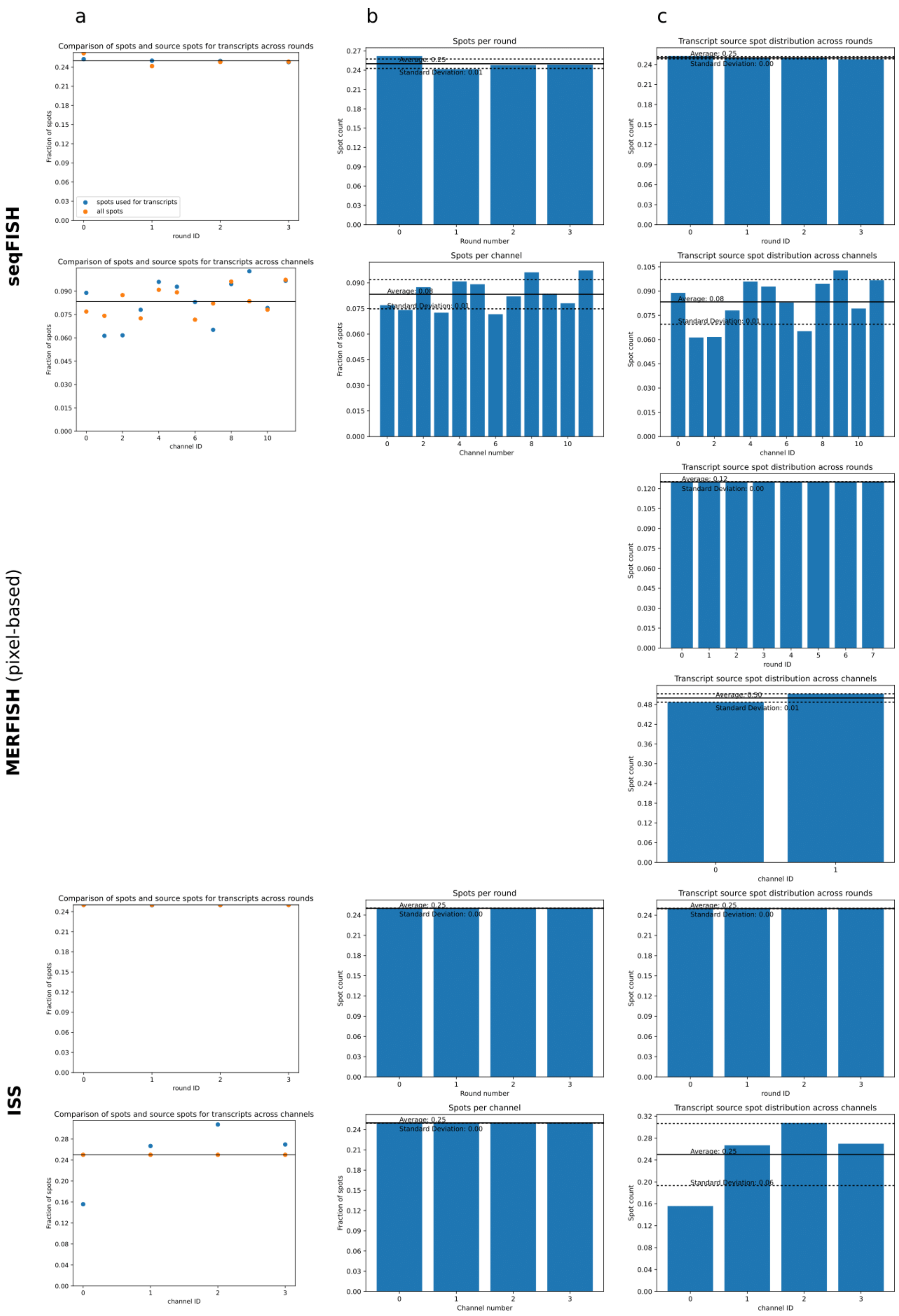
a
b
c
seqFISH
MERFISH (pixel-based)
ISS
Comparison of spots and source spots for transcripts across rounds
Fraction of spots
round ID
spots used for transcripts
all spots
Spots per round
Spot count
Round number
Average: 0.25
Standard Deviation: 0.01
Transcript source spot distribution across rounds
Average: 0.25
Standard Deviation: 0.00
round ID
Comparison of spots and source spots for transcripts across channels
channel ID
Spots per channel
Average: 0.08
Standard Deviation: 0.01
Channel number
Transcript source spot distribution across channels
Average: 0.08
Standard Deviation: 0.01
Transcript source spot distribution across rounds
Average: 0.12
Standard Deviation: 0.00
round ID
Transcript source spot distribution across channels
Average: 0.50
Standard Deviation: 0.01
channel ID
Comparison of spots and source spots for transcripts across rounds
round ID
Spots per round
Average: 0.25
Standard Deviation: 0.00
Round number
Transcript source spot distribution across rounds
Average: 0.25
Standard Deviation: 0.00
round ID
Comparison of spots and source spots for transcripts across channels
channel ID
Spots per channel
Average: 0.25
Standard Deviation: 0.00
Channel number
Transcript source spot distribution across channels
Average: 0.25
Standard Deviation: 0.06
channel ID

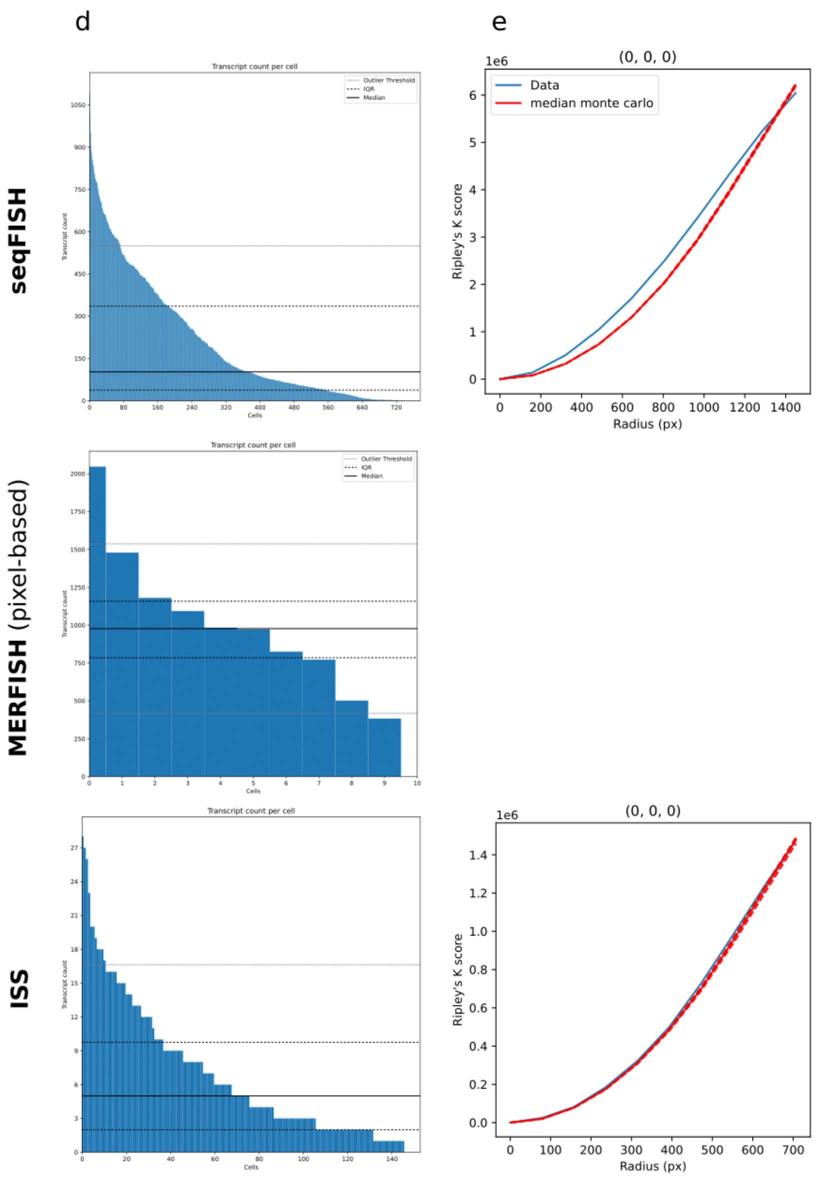


**Figure S3: Additional QC Metrics for each dataset.** Data is from the first FOV of each experiment, all plots are taken directly from pipeline output. Note that because MERFISH uses a pixel-based method, metrics that use spot data do not apply. A) Comparison of source spots for transcripts and all spots detected *(top)* across imaging channels and *(bottom)* across imaging rounds. B) Spot distribution *(top)* across rounds and *(bottom)* across channels. C) Transcript source spot distribution *(top)* across rounds and *(bottom)* across channels. D) Transcript count per cell. Outlier threshold is calculated as Median +/- 1.5*IQR E) Ripley's k score at increasing radius for spots, shown for the first z-slice of the first round and first channel. Solid line is median score under the null hypothesis (complete randomness) calculated by monte carlo method, dashed lines are 95% confidence interval calculated by monte carlo.

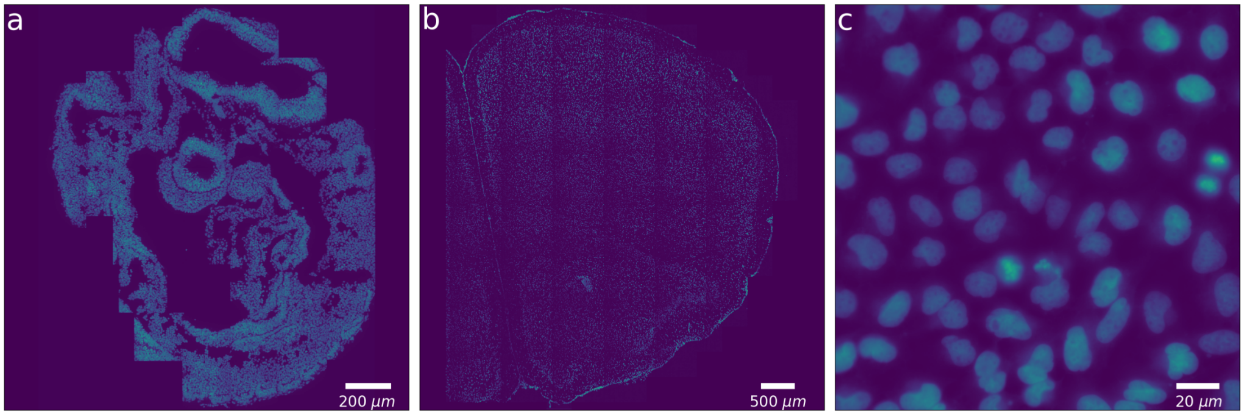


**Figure S4**: **DAPI channels of test datasets**
Full view of all assembled tiles for **a)** seqFISH (E8.5 mouse embryo), 351 genes measured across 40 tiles and 6 z-slices, **b)** targeted ISS (mouse coronal brain section), 50 genes measured across 190 tiles and 1 z-slice and **c)** a single example tile of the MERFISH (U2-OS Cell Culture), 130 genes measured across 495 tiles and 1-slice. Physical distance scales are included in each image.

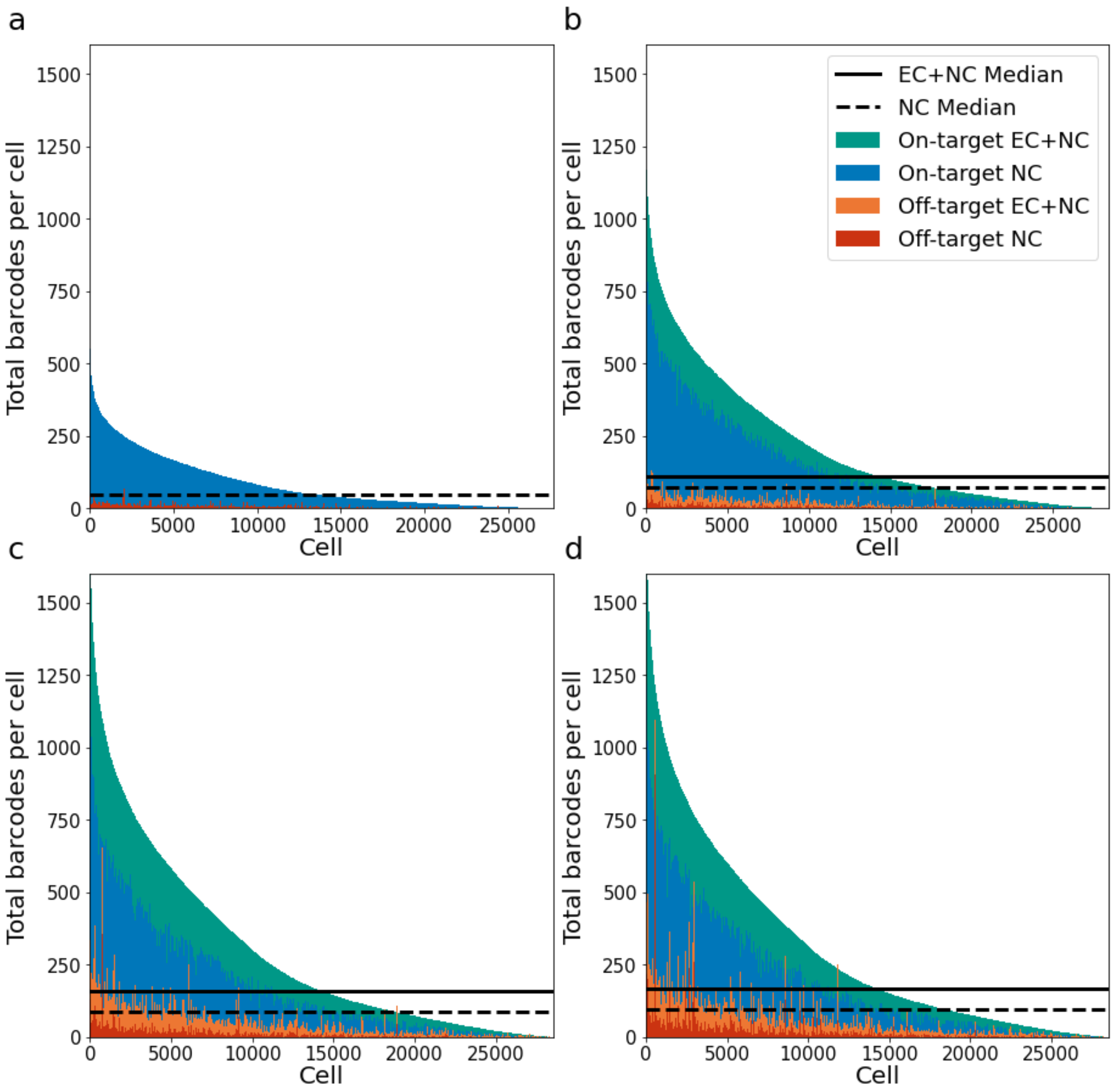


**Figure S5**: **False positive metric results for starfish and CheckAll decoders**
Comparison of performance for **a)** starfish NearestNeighbor decoder **b)** CheckAll decoder (high accuracy mode), **c)** CheckAll decoder (medium accuracy mode), and **d)** CheckAll decoder (low accuracy mode). Total counts of each barcode colored by barcode type. Error-corrected counts added to top of the non-corrected counts. EC = error-corrected, NC = non-corrected. Each column shows values for the same cell, ordered by their on-target NC+EC counts.

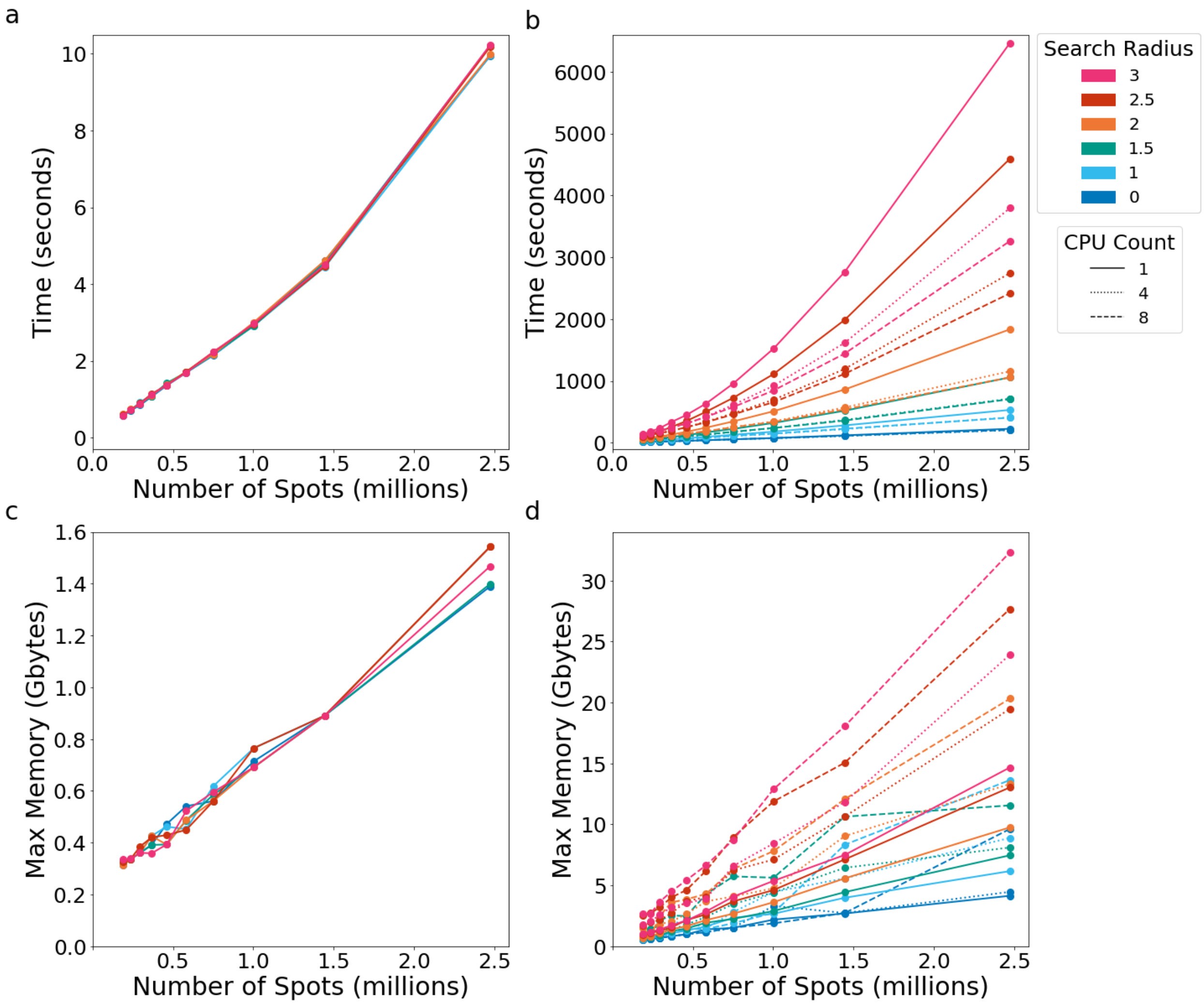


**Figure S6**: **Run time and memory benchmarks for starfish and CheckAll decoders**
Comparison of time benchmarks for **a)** starfish NearestNeighbor decoder, **b)** CheckAll decoder and memory benchmarks for **c)** starfish NearestNeighbor decoder, **d)** CheckAll decoder. Results shown for different numbers of input spots, search radii, and CPU's used for multiprocessing (CheckAll decoder only). Run on an AMD Ryzen 9 3900X (3.8 GHz base clock speed).

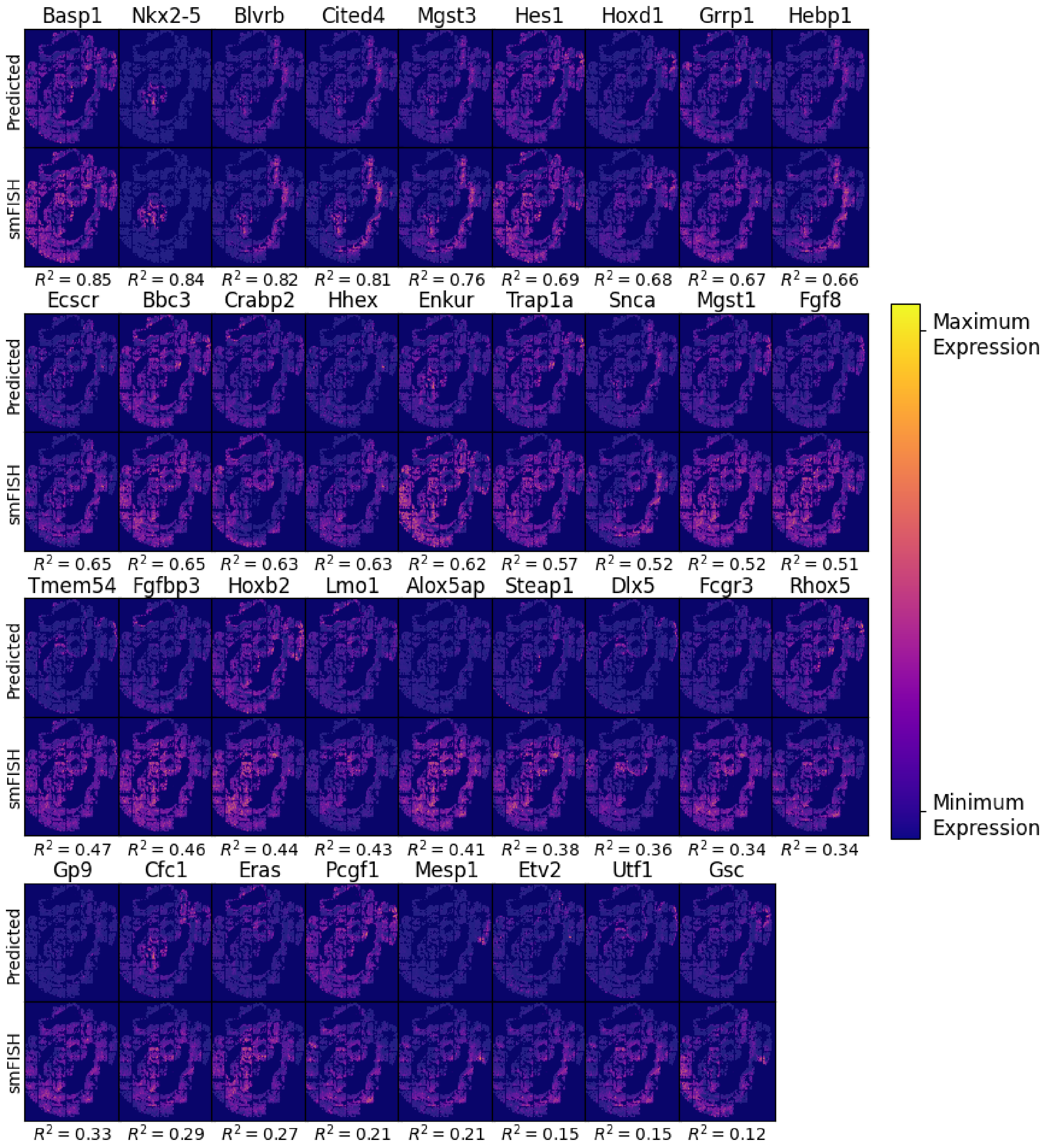


**Figure S7: Comparison of predicted and measured expression for all 36 smFISH genes**

For each gene, predicted expression by Tangram using seqFISH and scRNA-seq counts is shown on top and the measured expression by smFISH is shown on the bottom. The Pearson correlation of counts across all cells for each gene is printed below each and genes are ordered by the Pearson correlation. The gene ‘Ifng’ was measured by smFISH but is not shown here because it doesn’t appear in the scRNA-seq data used.

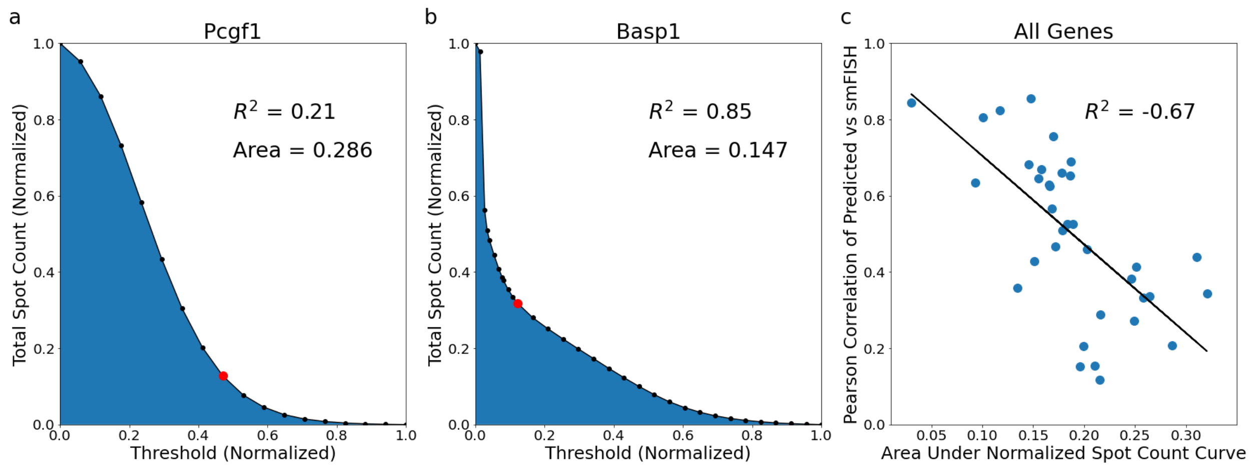


**Figure S8: Explanation of poorly performing genes for seqFISH external QC**
Normalized total spot vs threshold curves for smFISH images for **a)** Pcgf1 and **b)** Basp1. Printed $R^2$ is the Pearson correlation between predicted and smFISH counts for that gene while the area is the integral of the normalized total spot vs threshold curve. The red highlighted point indicates the elbow point of the curve that was used as the threshold value to obtain the printed Pearson correlation. **c)** Pearson correlation of predicted and smFISH counts vs the area under the normalized spot count curve for all genes. $R^2$ printed is the Pearson correlation between the Pearson correlation of predicted and smFISH counts and the area under the normalized spot count curve.

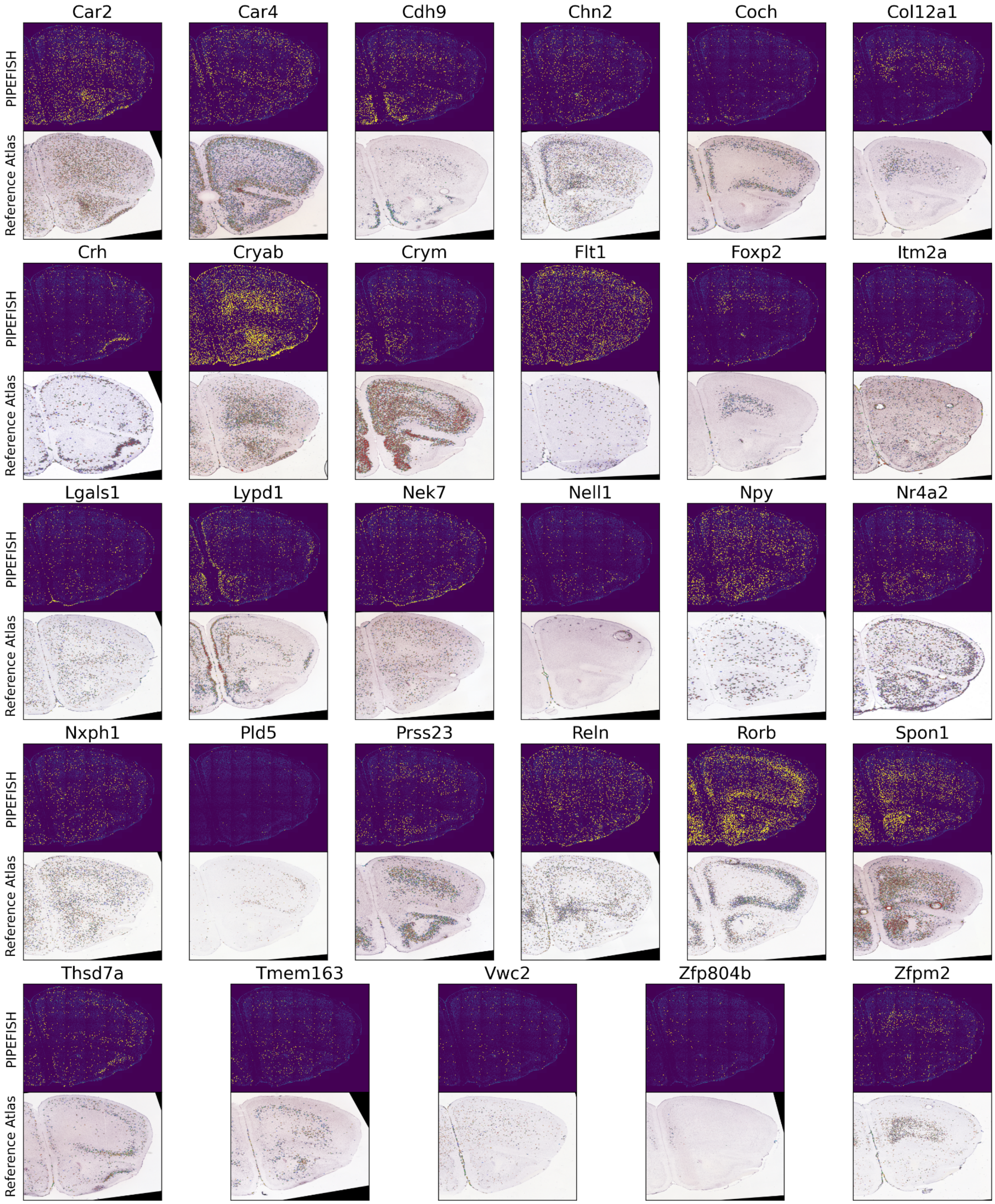


**Figure S9**: **Comparison of ISS pipeline expression with reference atlas**
All 29 additional genes found in the mouse brain ISS dataset with a coronal section reference in the Mouse Brain Atlas. In the top row: each yellow dot represents a transcript while the image underneath is the DAPI stain of the sample, in the bottom

row: blue dots represent low expression while more red dots represent higher expression (no color map provided by Allen Brain Atlas) while the image underneath is the Nissl stain of the sample.
